# Nitric oxide sensing by chlorophyll *a*

**DOI:** 10.1101/146563

**Authors:** Abhishek Bhattacharya, Pranjal Biswas, Puranjoy Kar, Piya Roychoudhury, Sankar Basu, Sanjay Ghosh, Kaustab Panda, Ruma Pal, Anjan Kr. Dasgupta

## Abstract

Nitric oxide (NO) acts as a signalling molecule that has direct and indirect regulatory roles in various functional processes in biology, though in plant kingdom its role is relatively unexplored. One reason for this is the fact that sensing of NO is always challenging. There are very few probes that can classify the different NO species. The present paper proposes a simple but straightforward way for sensing different NO species using chlorophyll, the source of inspiration being hemoglobin that serves as a NO sink in most mamalian system. The proposed method is able to classify NO from DETA-NONOate or (Z)-1-[N-(2-aminoethyl)-N-(2-ammonioethyl) amino] diazen-1-ium-1,2-diolate, nitrite, nitrate and S-nitrosothiol or SNO. This discrimination is carried out by chlorophyll *a* (chl a) at nano molar (nM) order of sensitivity and at 293K to 310K. Molecular docking reveals the differential binding behaviour of NO and SNO with chlorophyll, the predicted binding affinity matching with the experimental observation. Additional expreiments with diverse range of cyanobacteria reveals that apart from spectroscopic approach the proposed sensing module can be used in microscopic inspection of NO speices. Binding of NO is sensitive to tempertaure and static magnetic field. This provides additional support to the involvement of the porphyrin ring structure to the NO sensing process. This also broadens the scope of the sensing methods as hinted in the text.

## 1. Introduction

The importance of NO as a reactive free radical in biological environment is well documented [1]. However the role of NO in plant systems is relatively unexplored. A detailed understanding of the interaction mechanisms and knowledge in possible role of NO [2] may be of some interest. A number of extensive studies establish the significance of NO driven interferences in plant systems[3]. NO influences growth parameters, metabolism, cell differentiation, disease response, stress and nitrogen fixation etc.[4, 5, 6]. Again, detection, sensing and regulation of NO in photosynthetic organisms, cyanobacteria and plants are not clear[7].

A multiple of reductive and oxidative pathways are associated with NO biosynthesis in plant systems. Both enzymatic and non-enzymatic processes mediate NO production in plants, NO acting as an endogenous metabolite[8]. Additionally, NO production from L-arginine/NADPH mediated pathways by the activity of nitric oxide synthase (NOS)[9] enzyme generally involves a number of genes which further establish the enormous importance of such a small mediator molecule (NO) in plant systems[10, 11]. Exogenous conversion of NO from nitrite or other sources of NO further influence distinct physiological functions[12, 13, 14]. Although, extensive disparity and confusions regarding the activity of NO and limit of detection (LOD) for analytical systems arises due to unstable nature and high reactivity of NO[15].

Interestingly, NO is capable of effecting photosynthetic electron transition and initiate subsequent downstream cellular processes such as photophosphorylation [16, 17]. The macrocyclic core of the major photosynthetic pigment chla is coordinated to magnesium (*Mg*^2+^) as a central metal atom which is responsible for its unique optical, physical and biochemical properties [18, 19]. It is therefore logical to assume that the photosynthetic machinary would indirectly or directly ssense NO. This paper is primarily based on this rationale. A supportive argument could be the structural homology chlorophyll shares with hemoglobin that is a NO-sink in mammalian systems[20, 21, 22].

### 1.1. Reported methods for NO sensing

To address such problems regarding global significance and interest we investigate the interaction of both NO and SNO to the major pigment chl*a* at a cell free state and within cyanobacterial and algal photosynthetic membranes. Nitrates, nitrites, diolate or DETA-NONOate, Sodium nitroprusside or SNP and GSNO was also examined as other potent and reliable NO donors[23]. Additionally, NO reactivity depends on a number of factors and conditions such as temperature and oxygen pressure status of the micro-environment which readily interferes to the NO reactions. Finally, a molecular model has been proposed to study the interplay between the differential binding of NO or SNO to Chla. Importance of NO sensing is one of the major challenges with respect to the vast amount of interactions of this molecule in biological regulation [24, 25]. NO is generally detected as nitrite by griess chemistry (Peter Griess, 1858) and metalloprotein (HbO_2_-MetHb) assay systems[26, 27]. Additionally, a large number of NO detection tools are reported based on electrochemical analysis [28, 29], spectroscopic or fluorescence probes [30] such as fluorescein or DAF[31] and chemiluminescence methods[32]. Specialized and sophisticated instruments such as NO analyzers, electron paramagnetic resonance (EPR or ESR)[33] and direct contact based NO electrodes are available for the measurements of NO[34, 35]. However, these procedures are prone to a number of limitations such as specificity (non-specific and diffused signal), sensitivity (generally detect upto *μ*M to mM concentration ranges), donor selectivity[36] and source discrimination etc.[37, 38]. Photo-bleaching of the fluorophore, bio-compatibility, interference, cumbersome etc. are other issues that often hamper sensing [39, 40]. Disadvantages regarding direct contact based detection techniques or in case of penetrability also may be accounted contextually. There is again a huge need for photostable, non-toxic and selective dyes for NO imaging and bio sensing. Interestingly, this study might also have important implications in the development of novel NO sensing and monitoring technology and design of smart NO measurement green probes. NO monitoring methods at room temperatures along with photo active NO detection and static magnetic field (SMF) control over NO modulations at 77K further sensitize NO bio-sensing in the context of near infrared fluorescence (NIRF) emissions. NO detection and sensing by both cell free and membrane bound chl *a* had been studied.

### 1.2. Cyanobacterial photosynthesis and NO

A range of different unicellular *(Chlorococcum infusionum),* heterocystous *(Anabaena sphaerica)* and filamentous *(Leptolyngbya tenuis* and *Leptolyngbya valderiana)* algae and cyanobacteria had been examined to establish the universal role of chl*a* in NO biosensing. Heterocyst is a specialized nitrogen fixing cell distinct from the vegetative cell, found in some cyanobacteria and demarcated by an abundance of nitrogenase enzyme and absence of OEC[41, 42]. However, heterocyst formation depends on a number of specio-temporally regulated gene actions, nitrogen deprivation and 2-oxoglutarate (krebs cycle intermediate) etc. Additionally, heterocyst serve as a site for cellular differentiation in *Anabaena sphaerica.* Filamentous algae and cyanobacteria however differentiate by mechanical breaking or discontinuity in the filaments resulting in short vegetative strands which again self-differentiate to produce new filaments. Biotic and abiotic stress frequently influence filament breaking in *Leptolyngbya tenuis. Leptolyngbya valderiana* however exert high stability with regard to external stress response.

## 2. Materials and methods

Natural photosynthetic systems operate by a combination of different pigment molecules designated to perform coherent light dependent operations [43, 44]. Thus, separation and purifications of the component pigment molecules are a pre-requisite to study the effective role and activity of the individual components. Chl *a* was found to react to a diverse range of NO both at a cell free state and within photosynthetic cells.

### 2.1. Algal and cyanobacterial growth conditions

Unialgal cultures was purified and maintained in appropriate basal mediums at 20^o^C under 20-30*μ*mol photons *m*^−2^ *s*^−1^ illumination at a 16:8 hours of light:dark cycle. Bold basal media and cyanophycean medium was used for the growth and subculture maintenance of cyanobacterium *A. sphaerica* and eukaryotic green algae *C. infusionum* [45, 46]. Photosynthetic filamentous organisms were maintained at ASN III (artificial sea nutrient) culture media. SNO interactions and reactions were screened for a multiple of different unicellular, heterocystous and filamentous cyanobacteria and blue-green alga.

### 2.2. Isolation, separation and purification of chl a

Natural green pigment dye molecule chlorophyll was extracted from spinach ( *Spenacea oleracea)* leaves. Mesophyll tissue of the fresh green leaves was collected for extraction. The tissue was then subjected to freeze drying in presence of liquid nitrogen ( —196°C). The pigments were extracted in 2:1:1, methanol:petroleum ether:diethyl ether solvent system from the freeze-dried leaves. All the procedure was conducted at dark or dim light and at temperature regulated conditions. Vertical column (silica gel column chromatography, silica gel, SRL India, 100-200 mesh) and a planar thin-layer chromatography (TLC, DC kieselgel 60*F*_254_ Merck, Germany) surface was utilized to separate chl *a* from the the other analyte fractions using a volatile liquid/gas mixture of mobile phase[47]. Chl *a* is extremely sensitive to secondary degradations and are prone to photo bleaching and decay at a cell free state. The purity of chl *a* was verified by gradient HPLC analysis[48] (C-18 column, HPLC, Waters, USA). Major experimental data was validated by comparing to standard spinach chl*a* (C5753, Sigma-Aldrich, USA). Purified eluents were diluted in acetone and analyzed by UV-Vis and fluorescence spectrophotometry for the validation of chl *a* concentration (6*μ*g/ml)[49]. A pure chla specific spectral signature was evident with accurate position of B-band *(A_soret_)* and transition band (*A_Q_y__*) structures with distinct fluorescence emission signal at room temperatures and at cryogenic temperatures. Analytical grade PEG (81396, Sigma-Aldrich, USA), PVA (Sigma-Aldrich, USA), AOT (323586, Sigma-Aldrich, USA), MWNT and graphene was purchased and the noble metal nano particles were stabilized at a colloidal state following established methods[50, 51].

### 2.3. UV-Vis spectrophotometry

Absorbance measurements were conducted in an UV-Vis spectrophotometer (Thermo-Vision Evolution 300, USA). A xenon lamp source was used to illuminate the samples for the collection of absorbance data points. Spectral scan was conducted at a wavelength region between 300nm to 1000nm. Chl *a* exert an absorbance spectrum consisting of a soret band at 430nm and a Q_y_ transition band at 662nm (in acetone). Time dependent NO decay kinetics data was collected at fixed wavelength of 430nm. As the decay of NO is labile to temperature changes, all the experiments were performed at fixed temperatures controlled by a peltier system.

### 2.4. Fluorescence spectrophotometry

Steady state fluorescence measurements was performed in a PTI (Quantamaster^TM^40, USA). Excitation wavelength was set at 430nm with excitation and emission monochromators and emission was collected to a perpendicularly directed detector channels between 650nm to 750nm for chl*a.* The bandwidth was set at 5nm. Time kinetics data was collected with fixed excitation and emission wavelength set at 430nm and 665nm respectively. A synchronous fluorescence measurement was conducted at wavelength region of 350nm to 750nm with simultaneous excitation and emission scanning. All the experiments were performed at fixed temperatures as referred in the text. SNO treated samples were incubated for 30 minutes at 310K before measurements. All the experimental data was fitted to best fit theoretical models with coefficients having 95% confidence bounds using Origin 7 and Matlab 2014a software.

### 2.5. Cryogenic fluorescence measurements

Low temperature fluorescence emission was collected in a fluorescence spectrophotometer (Hitachi F-7000, Japan) equipped with a cryogenic low temperature measurement accessor (Model: 5J0-0112, Japan). Photosynthetic energy transfer dynamics (*P*_723_/*P*_700_), NO detection and spin coupled SNO sensing G-M switch was designed at 77K. The excitation light was provided from a xenon light source equipped with excitation monochromator set to 480nm (slit width = 5nm). Emission was scanned from 650 to 800 nm with emission monochromator (slit width = 10nm). The samples were diluted in presence of appropriate cryoprotectant glycerol (>60%) before freezing them prior to the low temperature measurements. Low concentration of free fluorophore order) was applied to avoid any interference due to self quenching or inner filter effects. Porphyrin specific excitation at 590nm was also implied at cryogenic temperatures[52] which results in multiple emission bands adjacent to the NIR region (data not shown).

### 2.6. Experimental NO reactants

GSNO is an endogenous S-nitrosothiol which is a source of bio-available NO[53, 54]. In an in-vitro temperature regulated liquid state GSNO releases maximum amount of purely dissolved NO. Intra-cellularly the amino acid glutathione[55] reacts to NO to form GSNO. Extracellularly GSNO decompose to generate NO and disulfide by a homolytic cleavage of S-N bonds[56, 57]. The following equation illustrates the decomposition of SNO either by thermal or photo lytic degradation or by transition metal-ion dependent breakdown into NO and thiyl radical RS^−^[58, 59, 57],

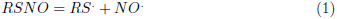

The *RS*. radical subsequently reacts to form di-sulfide (S-S) bonds. GSNO often catalyze S-nitrosation reactions[60, 61]. GSNO was prepared freshly at dark condition by mixing 1M aqueous solution of sodium nitrite *(NaNO_2_;* Merck, USA) with 1M solution of reduced glutathione (074011, GSH, SRL, India) in 1(N) HCl (HCl, Merck, USA) in 1:1 (v/v) ratio. GSNO formed was protected from light and placed on ice immediately. Further working dilutions (nM, *μ*M and mM ranges) were prepared freshly before experimental measurements. Nitrites (*NO*_2_^−^, 1 mM), nitrates (*NO*_3_^−^, 1 mM) (*N*_a_^2+^ or *K*+ salts), S-nitrosoglutathione (N4148, Sigma-Aldrich, USA), DETA-NONOate (82120, Cayman Chemicals, USA) and SNP (71750, SRL, India) were applied as other sources of NO. However, application of SNP results in nitrosonium ion (*NO*^+.^) and cyanide or *CN*^−^ as by-products which again acts as blockers of the photosynthetic electron transport chain[62]. Nitrites are again regarded as a potent source of NO[63]. Nitrates however undergo reduction reactions (NR enzymes) to generate nitrite and NO. Nitrites act as a vasodilator under hypoxic conditions and are regarded as potent diagnostic biomarkers[64, 65]. Metalloporphyrins are also considered as a precursor from which NO can be released from point irradiation by a photochemical de-nitrosylation process[66].

The impact of NO and SNO to chl *a* at a cell free state and at membrane bound conditions were analysed by incubation of the samples with referred amount of organic NO donors for 30 to 60 minutes at 37*^o^C* regulated by temperature and moisture controlled incubator at a closed condition to limit non-specific interferences. All the other reagents and solvents used are of analytical grade (*>* 99% purity) Sigma-Aldrich, USA, Merck, Germany and SRL, India products.

### 2.7. Purity testing of the NO donors

**Griess assay** Nitrite concentrations form the donor molecules were quantified using modified Griess reagent (G4410, Sigma-Aldrich, USA). Samples were mixed with 1× griess reagent and incubated for 30 to 60 minutes at 37^o^ C before measurements in uncoated 96-well plates with plate readers acquiring optical density at 540nm at 297K.

**DAF-2DA** The effectiveness of NO donors were verified and confirmed by 5mM diaminofluorescein diacetate (D2813, DAF-2DA, Sigma-Aldrich, USA) fluorescence emissions between 500-600nm with fixed excitation at 480nm.

### 2.8. Spin modulated quantum detection of SNO

An external SMF source of 500mT was primarily used to impart non-invasive spin perturbations to the fluorophores. A 10 minutes of magnetic (SMF) exposure was imparted to the samples at 277K before steady state and time kinetic photonic measurements at room temperature as well as at cryogenic temperatures. Previously, the 77K photosynthetic ratio (*P*_723_/*P*_689_) and membrane organizations was found to be responsive to SMF[67]. Notably, a 11nm red shift in the spectral position of the high energy (low λ) emission band of chl *a* results at 77K due to the solvent perturbation effects and hydrophobic self-interaction of the pigments. Contextually, a SNO sensing magneto-photonic G-M switching method had been designed at 77K. SNO intermediates was subjected to SMF at 277K at dark conditions before cryogenic measurements.

### 2.9. Fluorescence microscopy

Fluorescence imaging was conducted in a Zeiss anioscope a1 fluorescence microscope equipped with laser illumination source. Imaging was performed at 100 × magnification and in oil immersion. A phase contrast and fluorescence analysis of the photosynthetic organisms and subsequent SNO interactions were examined at room temperatures. Blue and red optical fiters and were utilized for the excitation and emission channels respectively for NO analysis.

### 2.10. Molecular docking

**Chlorophyll-A structure:** The Chlorophyll-A (chlA) structure was obtained from the PDB[68] file 3PL9 which is an X-ray structure of Spinach Minor Light-Harvesting Complex Cp29 solved at 2.80 *Ä*[69]. All hydrogen atoms were geometrically fixed to the chlorophyll-A molecule (residue identity: CLA, residue number: 604 in the PDB file) using the corresponding Autodock module and kollman partial charges were assigned to all atoms prior to docking[70]. The net charge of the molecule was considered +2 corresponding to the coordinated divalent cation, Mg(II).

### 2.11. Ligand structures

Structures of the reactive intermediates, namely, nitric oxide (NO) and S-nitrosothiols (SNO) were build by the popular structure drawing software Mar-vinSketch (https://www.chemaxon.com/products/marvin/marvinsketch), considering a partial triple bond and a double bond for N-O, respectively in the two molecules. All structures were optimized for their appropriate and accurate internal coordinates using the ‘clean in 3D’ option of the software. Marvin Sketch does not return any hydrogen atoms in the built structure which was necessary for the satisfaction of valencies and assignment of accurate charges to proceed for docking. Hence, the sulfhydryl hydrogen atom (−SH) was geometrically fixed externally by a previously validated program[71] using the technique of fourth atom fixation[72] with appropriate internal coordinates obtained from the literature[73]. Partial charges to the ligands were assigned using the classical Electronegativity Equalization Method (EEM) by the free online web server Atomic Charge Calculator[74].

### 2.12. Docking

The GUI interface of Autodock (version 4.2) was used for all the docking. The chlorophyll-A structure was fed as the macromolecule (receptor) and NO/HSNO as ligands. The grid boxes were visually set, centered on the receptor-centroid and enough large to cover all atoms of the whole receptor molecule giving rise to the dimensions of 40*Å* × 40*Å* × 40*Å*. No separate interaction spheres (centered on any preconceived active sites) were specified, so it was left on the docking algorithm entirely to determine the binding site of receptor-ligand pairs. Two independent ab-initio rigid body docking exercises were performed (one for each ligand) using a meta-heuristic GENETIC algorithm with its suggested default settings (population size 150, a maximum of 2500000 fitness evaluations, 27000 generations, gene mutation rate: 0.02, rate of cross-over: 0.8) for a run of 100 cycles. The resultant 100 conformers were then ranked according to binding energy (lower the better) and the best 15 models were selected for further analyses. The root mean square deviation of the ligand atomic coordinates among all conformations in the same cluster (say, all chlA-NO docked complexes) was also recorded and returned by autodock as cluster rmsd. Agreement in terms of these binding parameters among the selected docked conformations indicates convergence, meaning a definitive and stable mode of interaction. In such cases, the lowest binding energy conformer can be the best choice to convincingly represent the conformational ensemble and hence, the mode of interaction. Docked atomic models were displayed and analysed in Pymol (The PyMOL Molecular Graphics System, Version 1.8 Schrödinger, LLC).

## 3. Results

Tetrapyrrole ring currents of the major photosynthetic pigment chl *a* acts as a primary target of NO or SNO at a cell free state and at a membrane bound state. Hence, the action of chl *a* as an effective, robust and intrinsic NO sensing and monitoring tool is highly feasible. Although, a number of factors can influence NO reactivity, a liquid or solid state NO sensing method had been described at physiological temperatures. Comparable *R_f_* values were obtained for the column purified and TLC separated chl a. The rate of the relative migration of chl *a* was found to be 0.78 cm/min. *β*-carotene was separated with a *Rf* value of 0.963. Additionally, to understand the critical mechanism of how NO, SNO and other reactive nitrogen species effects photosynthesis[75, 76], a detailed understanding of the role of primary photon capturing unit chl *a* in balancing the electron and energy transition dynamics with contact to extracellular NO is necessary. At a cell free state, chl *a* in solution or at a thin film immobilized state or at membrane enclosed sub-cellular state can effectively replicate NO or SNO sensing at physiological temperatures.

### 3.1. Methods to probe NO reactivity

NO and SNO interact distinctly with chl *a* and exhibit reactant specific recognition and discrimination effects. SNO treatment to free chl *a* at a liquid state results in a drastic decay of both the soret and transition bands. About 19nm to 20nm blue shift (hypsochromic shift) of the soret maxima was noted accompanied with a profound degradation and vanishing of the *Qy* band at 37^o^C described in Fig. 1. SNO decompose gradually to liberate NO and di-sulfide adduct. The final product of the reaction, the chl-SNO intermediate decipher a novel absorption soret maxima at 410nm originated by the subsequent decay of the soret band of chl *a* at 430nm. Formation of SNO mediated nitrosated or nitrosylated macrocyclic superstructures with chl *a* are also plausible. Alternative to nitrosylation or nitrosation, SNO may lead to fluorescent adduct formation with the pyrrolic backbone of chl *a* at metal chelating sites (Fig. S1). GSNO concentration (nM) specific decay of both the soret and transition bands further justify intermediate complex formation. S-nitrosation or nitrosylation (NO incorporation) reactions of the fluorophore again may result in opening or closing of the chromophoric groups which in turn may impact (amplify or suppress) emissions. All the NO donors were calibrated by colorimetric griess assey (O.D at 540 nm) and DAF-2DA emissions for the quantification of nitrite (*NO*_2_^−^) and NO respectively at μM concentration ranges of detection limits. Nitrate (*NO*_3_^−^) was further reduced to nitrite (*NO*_2_^−^) with vanadium trichloride for diazotization reactions of griess assay (Fig. S2).

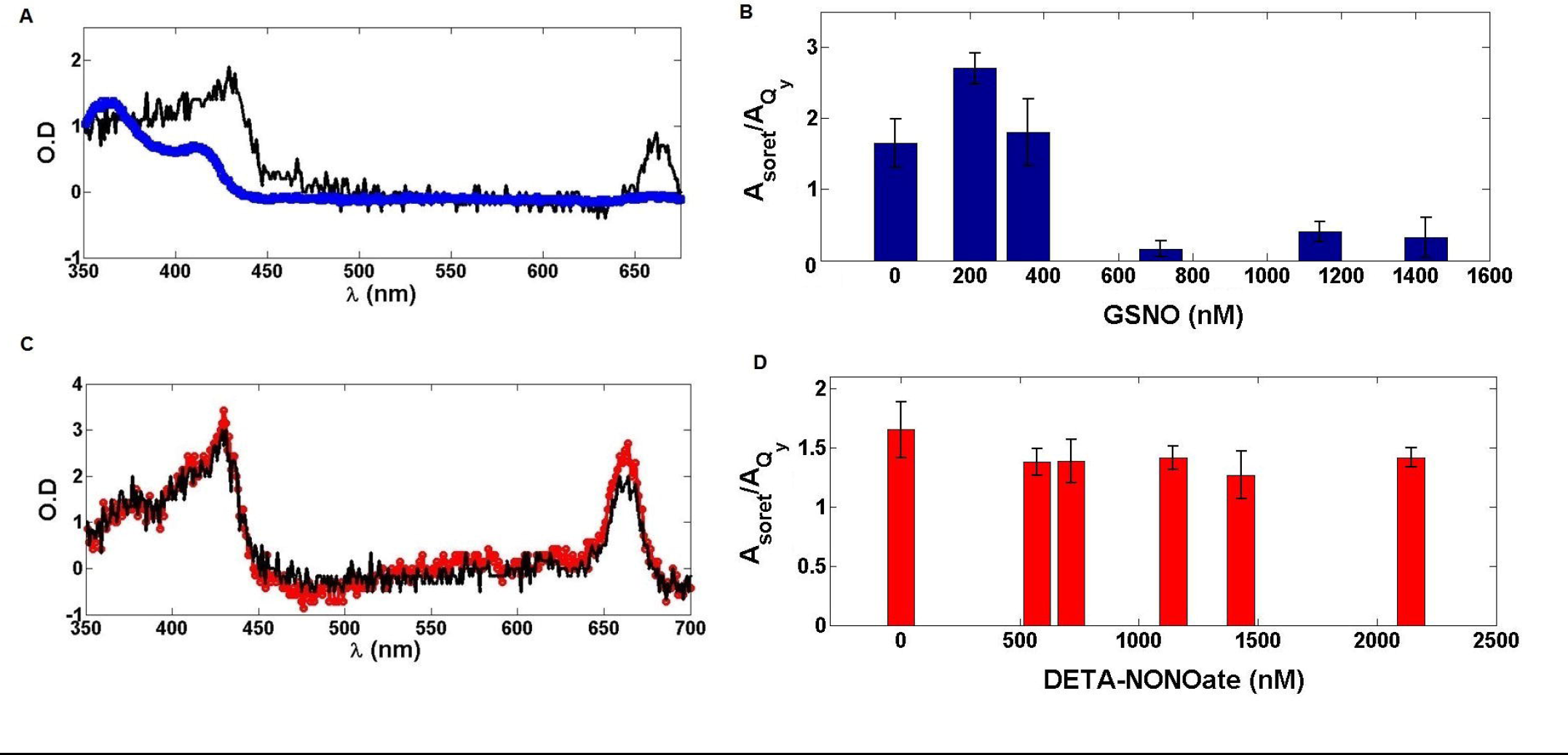

Alternatively, DETA-NONOate treatment results in a hyperchromic shift and a NO concentration dependent amplification of chl a. This gain was found to be consistent for both the soret and transition bands of chl *a.* SNO and NO mediated alterations of the absorbance (*A_soret_/A_q_y__*) ratio was described in Fig. 1 and Table 1. NO (DETA-NONOate) dose dependent spectral shifts again may frequently occur due to extensive π-π* self-interactions, non-degenerate excitonic interactions and polymerizations between structurally similar backbone rings. Transitions of the soret maxima (*O.D_max_*) further indicate towards the possibility of nitrosylated chl-NO intermediate complex formation while an increase in transition band (*A_Q_y__*) intensity generally refers to a gain in asymmetry of the π-electron distribution[77] and a gain in polarity of Chl-NO. Fluorescence emission channels again affirm distinct NO donor selectivity of the fluorophore at excited states with respect to exogenous NO and SNO corresponding to the alterations of the absorbance band structures. Fig. 2 describe the time kinetic fluorescence decay analysis at physiological temperatures which indicate higher NO release by SNO. The percentage rate of NO release at *t_50_* seconds from DETA-NONOate and GSNO was found to be 7.142s^−1^ and −73.04s^−1^ respectively (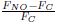 × 100, where *F_NO_* and *F_C_* denote NO/SNO treated emissions and un-treated emissions respectively). Notably, a negetive sign infer to the masking effect of SNO at 298K. Shifts in the transition band structures (Δ O.D) and the ground state decay kinetics again infer to the nature of the complex interactions and recognition effects of NO and SNO to chl a. Notably, emergence of an equilibrium extinction co-efficient (isosbestic point) at 422nm indicate reaction intermediate formation with SNO. Appearance of an isosbestic wavelength therefore confirms establishment of equilibrium and formation of complex intermediate products (1:1) between free fluorophore chl *a* and nitrosothiols. NO, nitrite or SNO specific ground state complex intermediate formation with the macrocyclic rings may adequately address the problem. Contextually, a number of immobilization substrates such as PEG, PVA, multi wall carbon nano structures, graphene, metal nano structures (gold, silver or iron nano particles), AOT (aerosol-OT) or alginate based hydrogel entrapment and chl polymer systems had been utilized for the stabilization of the pigments as solid state thin films for NO biosensing. Such bio-compatible, photo-thermally stable and re-usable bio-film systems can be utilized as a potent NO or SNO sensing device at room temperatures (Corresponding author Fig. S3). NO or SNO fabrication inturn prevents photo bleaching of chl *a* and imparts high photo-protection to the fluorescence intermediates. Interestingly, only a few nM concentration of NO donors are a pre-requisite to probe NO specific responses of the fluorophore. Surprisingly, a vectoral nature of NO and SNO detection and directional specificity and sensitivity had been noted for chl a. Additionally, the effects of NO to chl *a* was investigated under acidic and alkaline conditions and re-confirmed by checking the effects of the vehicles (NO dissolving medium) on chl a. The effects of dissolving medium such as polar and non-polar solvents, HCl, *H_2_SO_4_,* NaOH, *H_2_O, H_2_O_2_,* pH of the medium (100mM L-Asc, pH = 3 and NaOH, pH = 10), DMSO and PBS (pH, 7.4) on chl *a* had been examined. 1(N) HCl or NaOH results in expected degradation of chl *a* to one of its acid or alkali protoporphyrin derivative molecules which indicate a marked distinction in the spectral pattern with respect to the NO or SNO mediated changes of chl a. However, HCl lead to a quenching effect on chl fluorescence at room temperatures. NO/SNO mediated alterations of chl *a* can be further translated to a bio-luminescence platform under an UV-blue illumination at room temperature conditions. The effects of different exogenous nitric oxide sources on chl, the nature of reactant specific spectral shifts and NO detection and sensing had been illustrated in Table 1 and Table 2 respectively.

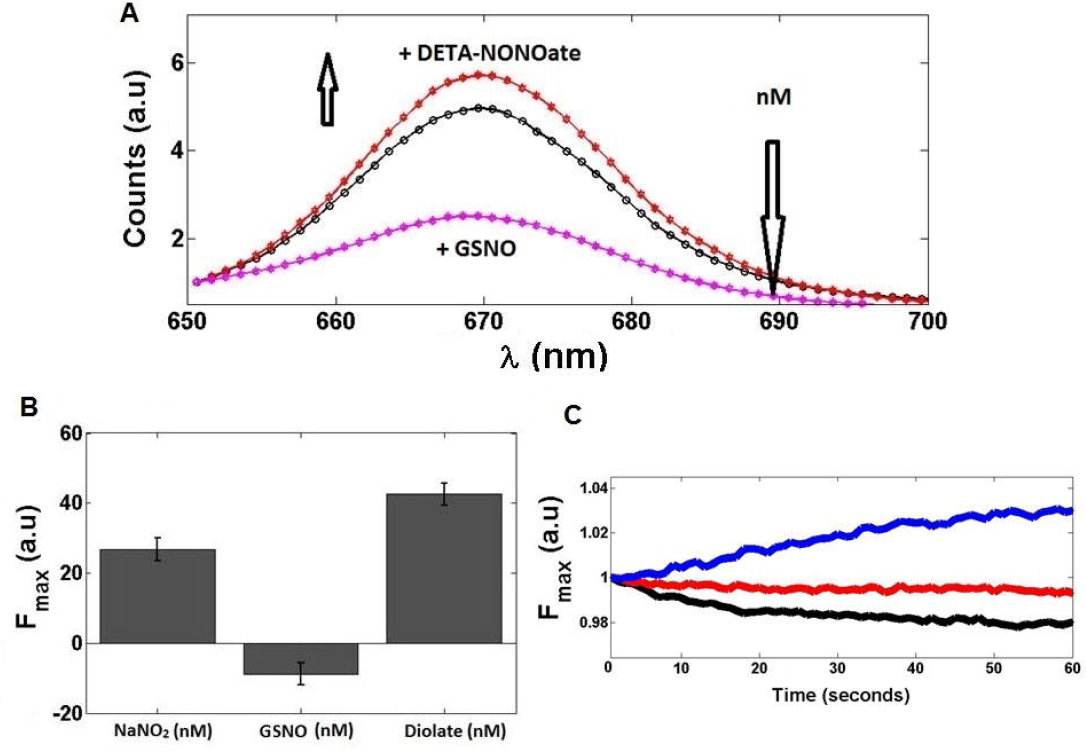

**Table 1.** Nature of spectral shift of Chla by NO donors. (−) represents not found.

| Sample | Nature of $\lambda$ shift | Soret band | $Q_y$ band | Isosbestic point |
| --- | --- | --- | --- | --- |
| Chla | (-) | 430nm | 662nm | (-) |
| Chla + GSNO | Hypsochromic shift (20nm) | 410nm | (-) | 422nm |
| Chla + DETA-NONOate | Hyperchromic shift | 430nm | 662nm | 440nm |
| Chla + $\text{NO}_2^-$ | Hypsochromic shift (99nm) | 331nm | (-) | 400nm |

**Table 2.** SNO switch. Fluorescence amplifications are represented by the ratio of the dual emission maxima of chla at 77K. (−) define not applied and (+) denote application of the factor in question.

| Chl$a$ (μM) | SNO (μM) | SMF | $P_{723}/P_{700}$ |
| --- | --- | --- | --- |
| 0.05 | (-) | (-) | 1.041 |
| 0.05 | (-) | (+) | 1.175 |
| 0.05 | 0.01 | (-) | 8.477 |
| 0.05 | 0.01 | (+) | 1.674 |
| 0.05 | 0.1 | (-) | 3.842 |
| 0.05 | 0.1 | (+) | 2.864 |
| 0.05 | 0.5 | (-) | 5.074 |
| 0.05 | 0.5 | (+) | 4.008 |
| 0.05 | 5 | (-) | 3.399 |
| 0.05 | 5 | (+) | 1.55 |
| 0.05 | 10 | (-) | 6.656 |
| 0.05 | 10 | (+) | 4.012 |

### 3.2. Characterizing the interactions by molecular docking

#### 3.2.1. Convergence of docking parameters

Large agreements were procured in the respective binding parameters (binding energy, cluster rmsd) among the best 15 conformations selected for each docking. In fact, the chlA-NO docked complexes returned practically identical binding energies (−1.84 ± 0.00 kcal/mol) for all 15 conformers with a cluster rms deviation of 0.018 ± 0.024 *Å*. On the other hand, for the docking of chlA-SNO, the corresponding values were −3.66 ± 0.01 kcal/mol and 0.30 ± 0.29 *Ä* respectively. The above results is a strong indicator of convergence and hence of a definite mode of interaction in both cases. The chlA-SNO docked conformations varied ever so slightly in their relative orientations (Fig. S4) whereas the chlA-NO docked conformations were practically identical. Hence, the lowest energy conformers (with a cluster rmsd of zero) were chosen as representative of the ensembles in either case.

#### 3.2.2. Mode of binding detected from key structural features

Note that the docking results are consistent and indicative of a definite mode of interaction without actually specifying any interaction spheres (or active sites) in the docking procedure (see section 2.12). The nature of both interactions were found to be electrostatic, represented by a close approach of the Mg(II) in the porphyrin ring and the O coming from NO / −SNO (Fig 3). The Mg-O bond length (O coming from −O=N) in chlA-NO was found to be 2.1 *A* falling in the same range to that of the MgN-pyroll coordinate bonds. Furthermore, the binding energies obtained for this electrostatic interaction (−1.84 kcal/mol) roughly falls into the same order of magnitudes to that of the hydrogen bond energies (1.4 kcal/mol)[78]. In comparison, the same Mg-O bond length in chlA-SNO was found to be much lower (1.7 *Å*), indicative of stronger bonding. The binding energy (−3.68 kcal/mol) was also found to be doubled in the later case. This difference in the strength of binding in the two interactions is consistent with the difference in their corresponding polarity, as reflected in the distribution of partial charges, in the two molecules. Surely, −SNO (N: −0.219, O: −0.327, S: 0.400, H: 0.145) is much more polar in comparison to NO (N: 0.131, O: −0.131) mediating a stronger electrostatic interaction with Mg(II).

The other interesting structural feature was the difference in the Mg-O=N bond angle in the two docked complexes: 160.7^o^ in NO whereas 115.5^o^ in −SNO. This means that NO was orientated in a close-to-linear manner to chlA making the docked complex relatively open to other exposures whereas the corresponding orientation in −SNO suggested significant bending, due to the incorporation of the bulky −SH group.

#### 3.2.3. Directionality in the binding modes of NO and SNO

Interestingly, NO and −SNO exhibited alternative directionality in their binding as revealed from the comparison of their best docked conformers (Fig. 3). Specifically, NO and −SNO binds to two opposite sides of the porphyrin ring-plane with respect to the location of the hydrophobic phytol tail. NO binds to the side where the phytol tail is bended and curving over the porphyrin ring whereas −SNO binds to the opposite ‘open’ side.

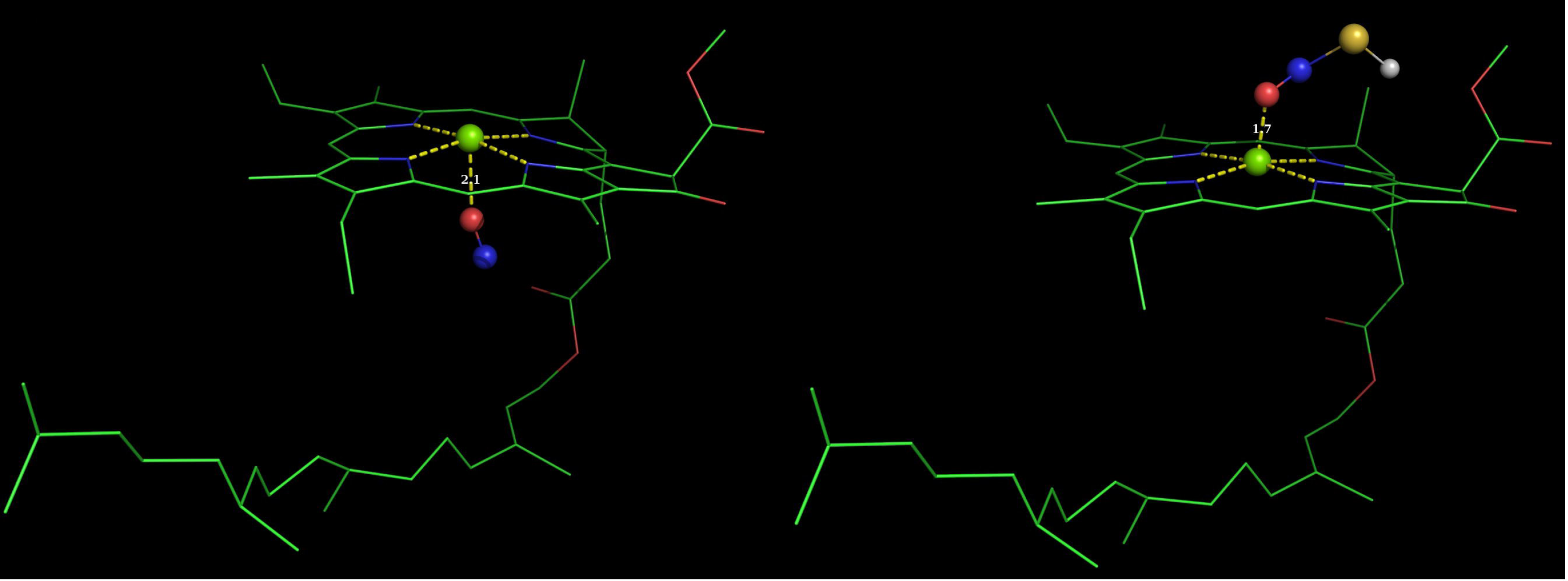

#### 3.2.4. Thermodynamic correlations with fluorescence experiments

The docking results are consistent and indicative of a definite mode of interaction. The nature of both interactions (NO and SNO) are electrostatic. The thermodynamic predictions from the docking study (i.e., binding energies) is such that the equilbrium constants for NO and SNO would be

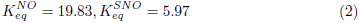

Interestingly, the observed fluoresent intensity ratio (which maybe approximated to the ratio of the NO:SNO equilibrium constant 19.83/5.97« 3.32) is of the same order 3.95 (see Fig. 3).

### 3.3. Magneto photonic Chl-SNO switch

At room temperature conditions, higher vibrational levels of the fluorophore become more populated due to thermal interference and agitation leading to broadening and masking of the fine electronic and vibrational informations. However, at liquid nitrogen temperatures (−196°C/77K) the thermal fluctuations render insignificant or negligible with less probability of solvent or medium collisions leading to a highly resolved spectral output. Contextually, a quantum sufficient NIRF detection method for NO (DETA-NONOate) and SNO had been developed. At cryogenic temperatures however the NO discriminatory behavior of chla render more prominent due to reduced or insignificant thermal noise and highly resolved band structures. At 77K, SNO favour down-hill energy transfer. DETA-NONOate however prefers an up-hill energy transfer described in Fig. 4. Notably, the weak spin coupled energy of SMF is also sensitive to high temperature destabilizations. SMF stabilize the *P*_723_/*P*_700_ ratio to greater than one (*P*_723_/*P*_700_>1) values with respect to control chla (where the *P*_723_/*P*_700_ ratio is equal to 1 denoting a thermodynamic equilibrium). Again, incubation of the fluorophore with SNO only results in even higher ratio values (*P*_723_/*P*_700_>>2). At cryogenic conditions extensive delocalized macrocyclic ring electronic currents preserves the optical and magnetic coherence and can be utilized as a spin selective indicator by prior exposure to a moderate intensity (500mT) SMF. As GSNO results in a concentration dependent drastic quenching of *P*_700_ with continuous enhancements of *P*_723_, a greater value of *P*_723_/*P*_700_ (>>2) was obtained each and every time with respect to equal amount of DETA-NONOate (*P*_723_/*P*_700_>1). Thus a range of increasing concentration gradient of GSNO was selected for the SMF coupled switching experiment. Fig. 5a describe a clear reversal and transition between the two states (G-state to M-state) for the fluorophore. Interestingly, SNO reactivity had been probed exploiting the spin chemistry of the extensive delocalized π-electronic currents of chl *a* with respect to external SMF at 77K. The *P*_723_ band exert higher response to SNO if considered separately. Fig. 5b illustrate an alternate nature of fluorescence switching by the *P*_700_ and *P*_723_ nm bands with respect to GSNO (G) and SMF (M). SNO treated chl *a* (G-state, denoted by +1) upon magnetic (SMF) exposure results in a spin coupled fluorescence reversal and a decrease in the *P*_723_/*P*_700_ ratio (M-state, denoted by −1). SNO complementarity by SMF infer to the development of novel and effective methods to probe NO and SNO. Design of an optically detectable magnetic switch for SNO detection had been described in Fig 5c. Such SMF reversibility of the SNO treated chla originate from reactant specific net magnetic moments and free energy of the reaction products. The operation of the switch is guided by a reactant specific spin selectivity and a magnetic reversal pathway (Table 2).

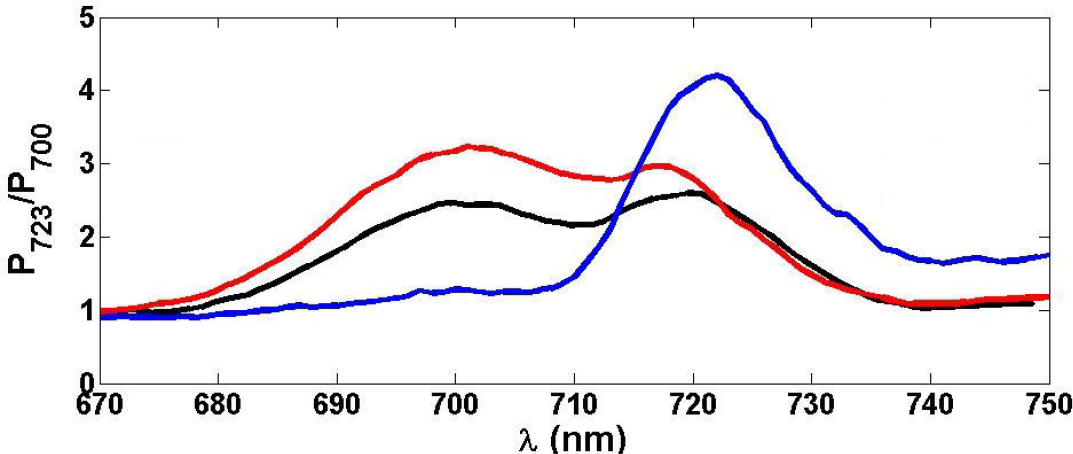

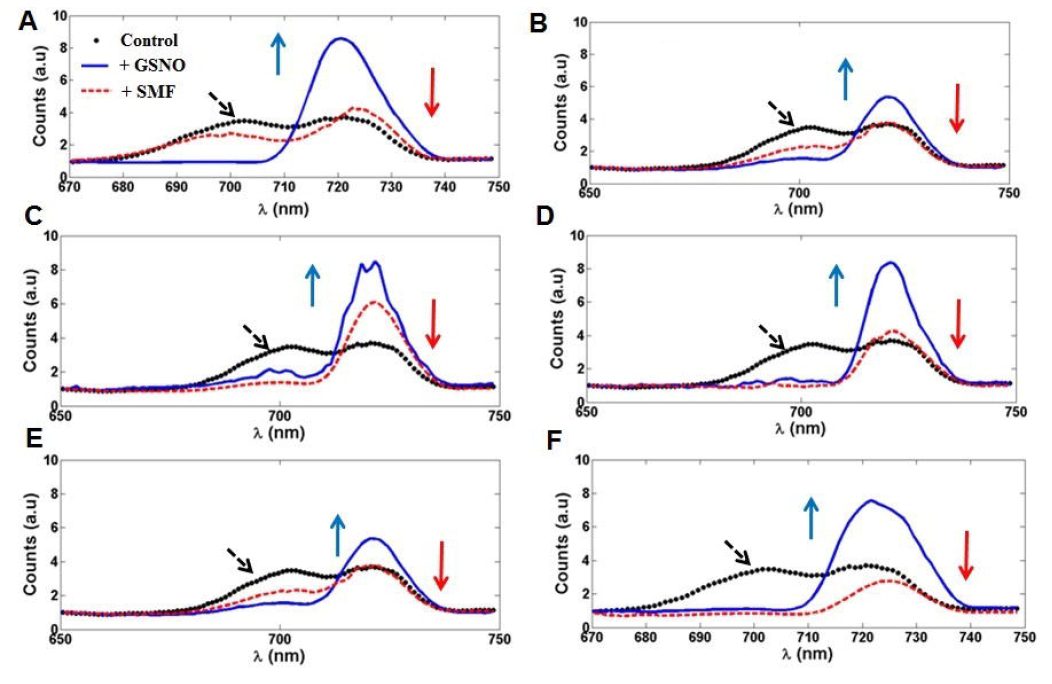

NO or SNO react to chl*a* to alter the energy transfer dynamics, free-energy and the net magnetic moment of the fluorophore. A non-invasive control over the SNO mediated reaction products was established by using an external SMF which at such conditions tend to preserve the free-energy and equilibrium and functions by regulating the net spin magnetic moments of the products. A bi-modal magneto photonic G-M switch had been designed for SNO.

### 3.4. Cyanobacterial mapping of SNO

SNO dependent drastic quenching of the major photosynthetic pigment chla specific auto-fluorescence was evident for all the algal and cyanobacterial cell types described in Fig. 6a. Additionally, a four to ten times less intensity in fluorescence emissions was noted after SNO treatment and incubation which suggests reaction intermediate formation in-vivo. A distinct shift in the fluorescence emission histogram was noted for the unicellular *C. infusionum.* A membrane level fluorescence analysis of *C. infusionum* after SNO application (200μM final concentration) and a nM order of SNO dose response had been studied at 310K (Fig. S5). *A. sphaerica* however exert marked and distinctive cellular alterations after SNO incubation. A change in the mode of heterocyst activity and cellular differentiation had been observed. Highly effected heterocysts frequently switch to an amplified cellular differentiation mode to form distinct and separate vegetative cells. Additionally, SNO treatment results in an elevation of the intra-cellular micro-oxic conditions which limits oxygen supply, protects nitrogenase and increase the sub-cellular NO concentrations in the heterocysts. Rise in the intra-cellular NO level deliberately accelerates heterocyst transformation to vegetative cells for enhanced differentiation, multiplication and generation of separate individuals of *A. sphaerica* cells (Fig. S6). Non-heterocystous and filamentous cyanobacteria *L. tenuis* again exhibit high activity of the stress generated tiny vegetative fragments for higher multiplication and generation of new filaments by SNO. *L. valderiana* however exhibit highest level of cellular nitrite synthesis compared to the other unicellular, heterocystous and filamentous cyanobacteria described in Fig. 6b. Persistent quenching of chla fluorescence emission by SNO was again verified for *L. valderiana.*

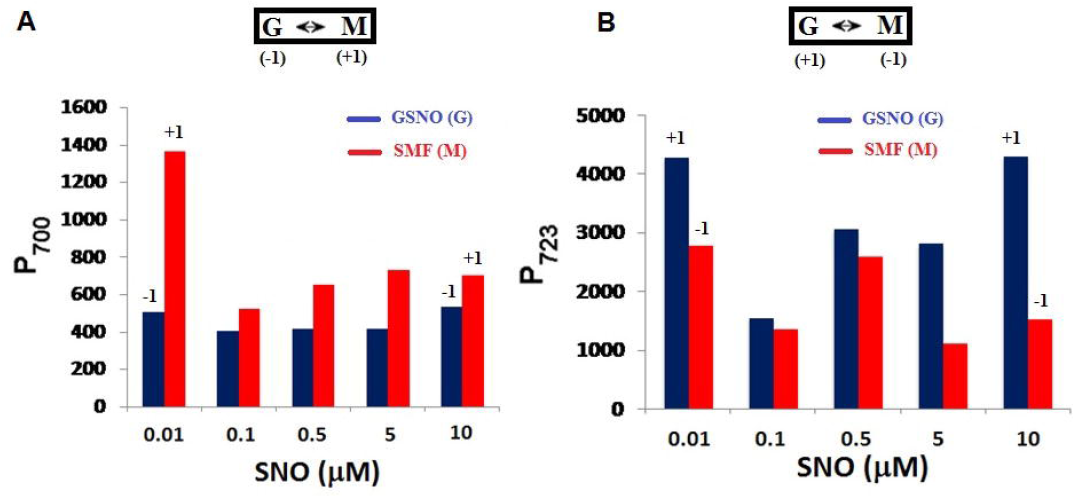

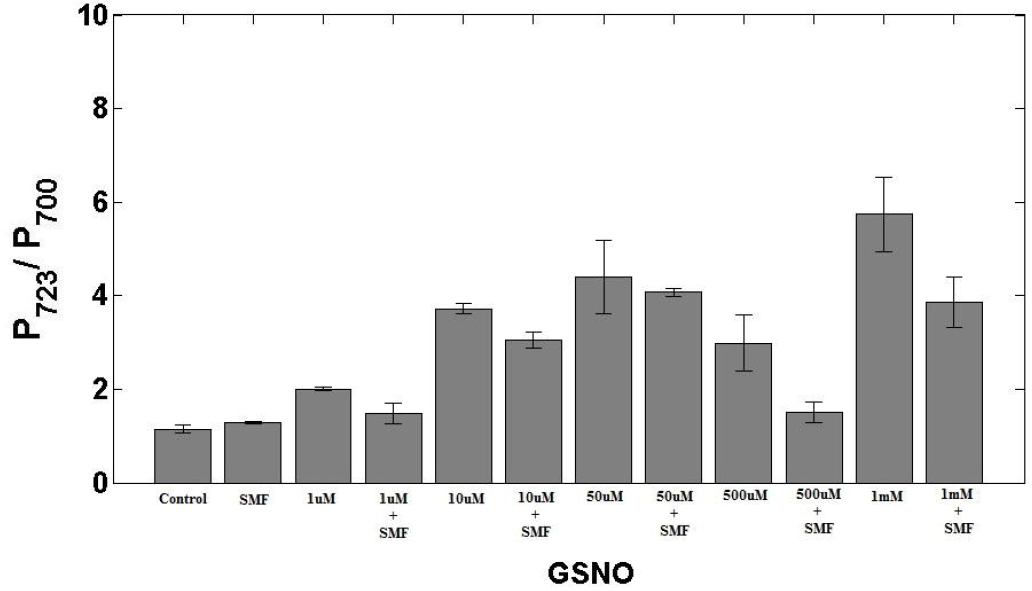

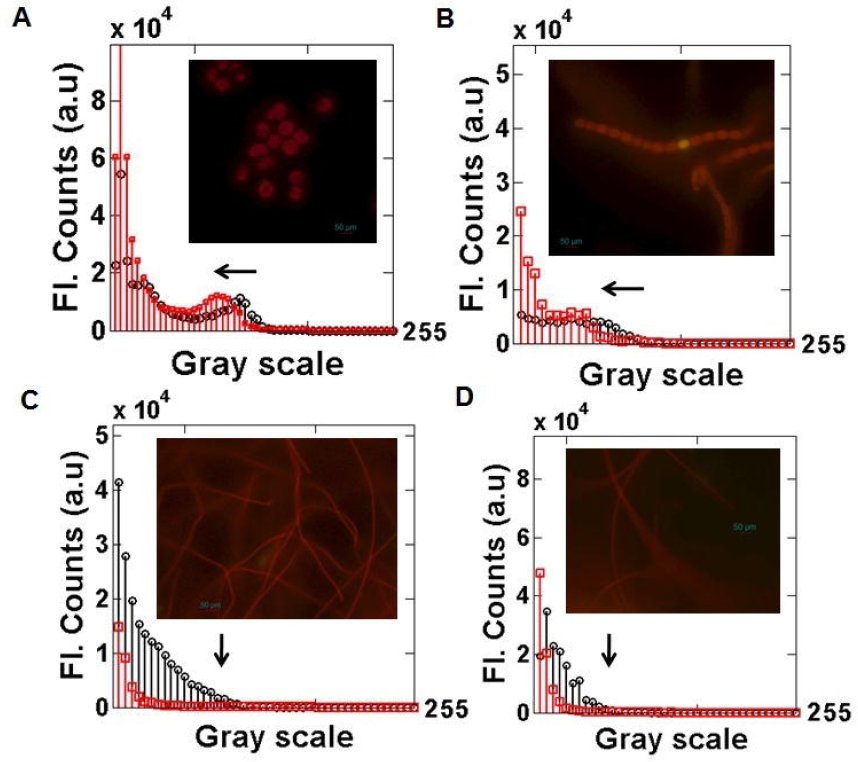

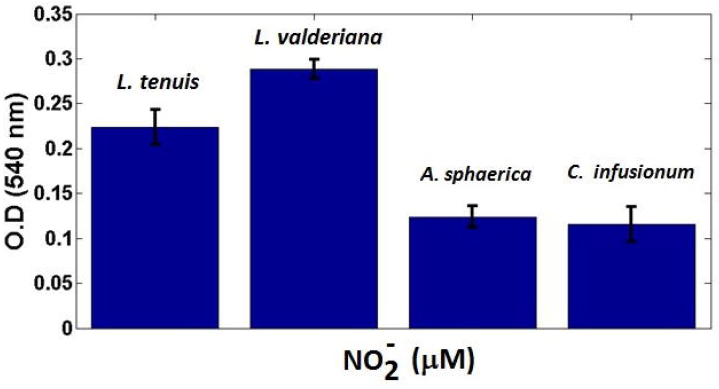

## 4. Discussions

Major photosynthetic pigment chla was reported to effectively interact to NO at a donor specific manner to identify and discriminate between SNO and other NO sources such as DETA-NONOate. Notably, a nM order of sensitivity in the detection limit had been observed. Such a low level of sensing of NO and SNO is a matter of serious interest. Interestingly, NO sensing was noted for both in-vivo membrane bound chl*a* and at an in-vitro cell free state at physiological temperatures. Giant spectral frame shifts of the intrinsic band structures of the fluorophore and NO dose dependent response of the same infer to metal chelations, nitrosylation, pigment polymerization, macro-cyclic fabrications and nitrosation reactions at a reactant specific way. The proposed methods mainly rely on easily accessible, inexpensive and replicable spectrophotometric, fluorescence and imaging methods. Alternative to this, a MS (mass spectrometry) based analysis would be useful to decipher the reaction intermediates and products at a molecular level and is a subject matter of future prospects of the work. Prospects for the design of novel memory enabled quantum nano bio materials, photo sensitized NO sensing, plant health monitoring, pesticide screening, disease marker related problems and material characterizations are the relevant issues that can be addressed utilizing the current method.

Additionally, algal and cyanobacterial screening of SNO at an interface of in-vivo membrane bound chl*a* correlate to the subsequent functional and physiological reactivity of the photosynthesizing organisms to SNO. Specialized nitrogen fixing cells exert a SNO dependent cellular differentiation and dissolution of the heterocysts to transform into separate vegetative filaments. A shift of the fluorescence emission maxima along with a drastically quenched and supressed intrinsic fluorescence emissions of chla was noted upon GSNO incubations. SMF complementarity and SNO reversibility of the cryogenic emission ratio (*P*_723_/*P*_700_) further supports chl-SNO intermediate stabilizations and a noninvasive control over the accessibility of the chromophoric groups. Classical perspective for such reactant specific magnetic interactions substantiate to a field mediated rotation of charged particles. While quantum angular momentum, spin and net magnetic moment corroborate to the optical memory, spin coupling and SNO switch. Lastly, chl*a* may act as a sub-cellular reservoir for NO by converting into an analogue of heme which share a common precursor of biosynthesis.

## 5. Conclusions

Quantum transitions of chla upon NO incubation infer to a reactant specific chromophoric accessibility and orientation, selectivity, specificity, biasness and sensitivity depending on specific intermediate formation at physiological temperatures. Compared to the neural circuits (quantum) of the brain [79], chla also interact exclusively to distinct NO donors or sources. Interestingly, the proposed method detects NO at nM concentration ranges and corroborate the possibility of NO derived magnetic memory in chl*a.*

NO sensors are classically intended to be bound on the principles that obey the classical analytical approach in which a linear response is expected. Sensing of this extremely complex system can also be approached using pattern recognition in which changes in the specific spectral pattern signify to the presence of a particular NO donor or SNO and its accurate distinction by artificial intelligence. Contextually, development of a new sensor class had been illustrated suitable for nM detection limit for the universal regulatory molecule NO. Lastly, the observations may throw some light on a hitherto unexplored domain for the intrinsic NO sensing mechanisms in photosynthetic, plant and other biological systems.

## Abbreviations

Chla: Chlorophylla
NO: Nitric oxide
*NO*_2_^−^: Nitrite
*NO*_3_^−^: Nitrate
*NO*_x_: Nitrogen oxides
GSNO/SNO: S-nitroso-glutathione
DETA-NONOate/ Diolate: (Z)-1-[N-(2-aminoethyl)-N-(2-ammonioethyl) amino] diazen-1-ium-1,2-diolate
SNP: Sodium nitroprusside
NR: Nitrate/Nitrite reductase
TLC: Thin layer chromatography
*R_f_*: Retardation/Retention factor
HPLC: High pressure liquid chromatography
*HbO*_2_: Oxyhemoglobin
MetHb: Methe-moglobin
*HCO*_3_^−^: Bicarbonate
GSH: L-glutathione reduced
CN^−^: Cyanide
NADPH: Nicotinamide adenine dinucleotide phosphate
PBS: phosphate buffer saline
DAF-2DA: Diaminofluorescein diacetate
PEG: Polyethylene glycol
PVA: Polyvinyl alcohol
AOT: Bis(2-ethylhexyl) sulfosuccinate sodium salt
nM: Nano molar
*μ*M: Micro molar
nm: Nanometer
O.D: Optical density
K: kelvin
mT: Milli-tesla
SMF: Static magnetic field
G: gauss
Oe: Oersted
NIRF: Near infra-red fluorescence
OEC: Oxygen evolving complex

## Acknowledgements

The authors like to thank Department of Biotechnology, Govt. of India (BT/PR3957/NNT/28/659/2013) for providing funds. We thank DBT-CU-IPLS and CRNN-CU for providing high end infrastructural facilities. We also thank Mr. Souradipta Ganguly and Mr. Sourav Kumar Patra for their assistance regarding NO testing. The work was partly supported by the Department of Science and Technology Science and Engineering Research Board (DST-SERB research grant PDF/2015/001079). We show our gratitude to The Center of Excellence in Systems Biology and Biomedical Engineering (CoE), University of Calcutta.

